# Multiplexed photo-activation of mRNA with single-cell resolution

**DOI:** 10.1101/2020.03.19.999631

**Authors:** Dongyang Zhang, Shuaijiang Jin, Xijun Piao, Neal K. Devaraj

## Abstract

We demonstrate sequential optical activation of two types of mRNAs in the same mammalian cell through the sequential photocleavage of small molecule caging groups (‘photo-cages’) tethered to the 5′ untranslated region (5′-UTR) of an mRNA. Synthetic ‘photo-cages’ were conjugated onto target mRNA using RNA-TAG, an enzymatic site-specific RNA modification technique. Translation of mRNA was severely reduced upon conjugation of the ‘photo-cages’ onto the 5′-UTR. However, subsequent photo-release of the ‘cages’ from the mRNA transcript triggered activation of translation with single-cell spatiotemporal resolution. To achieve sequential photo-activation of two mRNAs in the same cell, we synthesized a pair of ‘photo-cages’ which can be selectively cleaved from mRNA upon photo-irradiation with different wavelengths of light. Sequential photo-activation of two mRNAs enabled precise optical control of translation of two unique transcripts. We believe that this modular approach to precisely and rapidly control gene expression will serve as a powerful tool in future biological studies that require controlling translation of multiple transcripts with high spatiotemporal resolution.

## INTRODUCTION

The ability to precisely control gene expression is important for a wide range of applications in basic biological research, genetics, as well as gene therapies. [1-2] Stimuli-responsive control in confined time and space is enabled by inducible gene expression systems. For instance, small-molecule inducers, such as doxycycline, have been widely used in regulating synthetic gene circuits. [3] However, chemical inducers have slow diffusion rates, potential toxicity, and off-target effects on living systems, limiting their utility when high spatiotemporal resolution is required. [4] In contrast, optogenetic approaches utilize light as an external stimulus to regulate cellular processes. [5] Light irradiation is convenient to apply to biological samples such as live cells and organisms. The adverse effects of light-irradiation on living systems can be minimized by optimizing the wavelength and irradiation period of the light source. Perhaps most importantly, light can be applied with high spatial-temporal resolution, offering fast activation and deactivation dynamics. [6-7]

There has been an extensive body of research aimed at developing optogenetic tools to regulate gene expression. In a common strategy, a light-sensitive protecting group (‘photo-cage’) is chemically installed on a biologically relevant target (e.g. metabolites, oligonucleotides, or proteins) rendering the substrate inactive. Subsequently, light irradiation triggers the release of the ‘photo-cage’ from the target biological molecule, restoring its cellular activity. [8-10] For instance, optogenetic approaches often involve the installation of light-responsive protein domains or amino acids to achieve photo-chemical manipulation of proteins such as nucleases, proteases and transcription factors. [11-16] Caged oligonucleotides have also been used to control gene expression at the level of transcription or translation. [17-24] By conjugating ‘photo-cages’ onto nucleotide bases, hybridization between the antisense oligonucleotide and the target DNA/RNA can be controlled precisely using light. While most studies have focused on optogenetic control of transcription, there are advantages to methods that optically control gene expression at the level of translation. [25-29, 41] Since mRNA can be processed by cellular translation machinery immediately upon cytoplasm entry, controlling gene expression through direct manipulation of mRNA provides more rapid changes in cellular protein concentration compared to the regulation of transcription. [30] Moreover, *in-vitro* transcribed mRNA (IVT-mRNA) is only transiently active in cytosol and is completely degraded via cellular metabolism, typically within 24 hours. Thus, unlike the use of plasmid DNA or viral vectors which may be integrated into the cellular genome, mRNA-based gene expression regulation systems do not pose the risk of insertional mutagenesis. [31] Thus, a technique that enables optogenetic manipulation of translation would benefit from the high spatial-temporal resolution inherent to optical control as well as fast dynamics of mRNA processing.

Previously, we demonstrated a technique which enabled precise optical control of mRNA translation through the conjugation of light-sensitive ‘cages’ onto IVT-mRNA. [32] Laser irradiation (405 nm) on live cells removed the ‘photo-cages’ from cytoplasmic mRNA, subsequently activating translation with single cell precision. We speculated we could significantly expand this tool by enabling sequential photo-activation of two mRNAs within the same cell. Here, leveraging the multiplex capability of our mRNA caging/uncaging platform, we describe a sequentially light-activated translation regulatory system that utilizes two ‘photo-cages’ to cage two types of mRNAs. Irradiation with longer wavelength light (456-488 nm) activates one mRNA, while subsequent irradiation with shorter wavelength light (365-405 nm) activates the other mRNA. This multiplexed gene expression regulatory system provides a high degree of flexibility and shows potential for enabling the study of multiple regulatory genes with high spatial-temporal resolution.

## RESULTS AND DISCUSSION

To allow for site-specific and covalent conjugation of small molecule effectors onto target mRNA, we previously developed a technology named RNA transglycosylation at guanosine (RNA-TAG), which utilizes a bacterial tRNA guanine transglycosylase (TGT) to exchange a guanine nucleobase within a specific 17-nucleotide RNA stem-loop structure (‘Tag’) with synthetic enzyme substrate analogs (Figure S1, Figure 1A). [33-34] In *E. Coli*, tRNAs carrying Asn, Asp, His and Tyr are post-transcriptionally modified by TGT, which exchanges the guanine at the wobble position of the anticodon loop of the tRNA with the enzyme’s natural substrate pre-queuosine1 (preQ_1_). [35-36] Recognition of the RNA substrate by TGT does not require the full sequence of tRNA. Instead, a minimal 17-nucleotide RNA stem-loop from the anticodon loop of the tRNA (the 17-nucleotide ‘Tag’ sequence) is sufficient to promote TGT recognition and labeling. [37-38] TGT also accepts a wide range of synthetic preQ_1_ derivatives as small molecule substrates. By genetically inserting the ‘Tag’ sequence into an RNA of interest, we previously demonstrated TGT labeling is able to covalently conjugate a variety of functional small molecules, such as fluorophores and affinity tags, site-specifically onto the target RNA. [33-34] The versatility of this RNA modifying platform enabled us to adapt this technique to regulate translation, through conjugation of small-molecule effectors directly onto an mRNA transcript.

**Figure 1.**
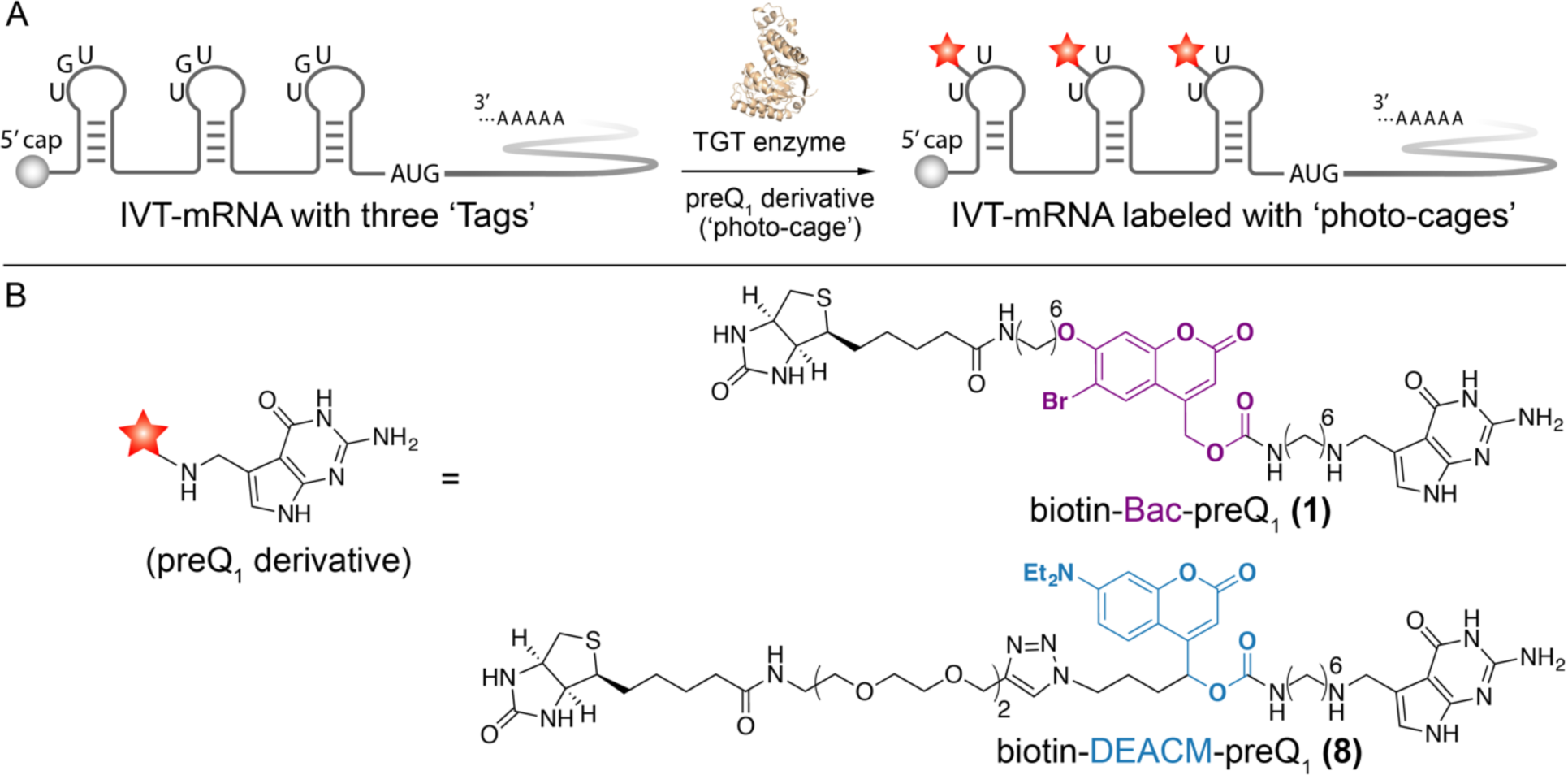
mRNA photo-caging using the RNA-TAG technique. A) To facilitate TGT enzymatic labeling, three enzyme recognition sequences, ‘Tags’, are genetically inserted along the 5′-UTR of an IVT-mRNA. Subsequent conjugation of the ‘photo-cages’ severely reduces mRNA translation activity. B) Chemical structures of two sequentially activable preQ_1_ derivatives (‘photo-cages’), biotin-Bac-preQ_1_ and biotin-DEACM-preQ_1_.

To achieve optical-control of mRNA translation, we covalently conjugated the synthetic ‘photo-cage’, biotin-Bac-preQ_1_ **(1)**, at three different locations along the 5′-UTR of a mature IVT-mRNA (Figure 1). The first conjugation site was located adjacent to the 5′-cap of the mRNA, specifically, at the 11th base after the 5′-cap. The second conjugation site was located in the middle of the 5′-UTR. The third conjugation was located at the 11th base upstream of the AUG start codon (Figure 1A). To determine the effect of such conjugation on translation, both the labeled and unlabeled IVT-mRNAs were delivered into mammalian cells through transient transfection. As a result, translation efficiency of the labeled IVT-mRNA was severely reduced to approximately 10% of the activity relative to the unlabeled IVT-mRNA. [32] We hypothesized that this phenomenon is due to the steric hindrance of these bulky ‘photo-cages’ conjugated at the 5′-UTR, which is where translation initiation takes place. Upon irradiation with 405 nm laser light, the Bac linker was photo-cleaved, leaving a minimal amine residue on the mRNA. As a result, translation activity of the IVT-mRNA was recovered after photo-uncaging, demonstrating successful photo-activation of gene expression. Importantly, mRNA translation was only observed in laser irradiated cells, and not in adjacent cells, demonstrating single-cell resolution of activation.

**Scheme 1.**
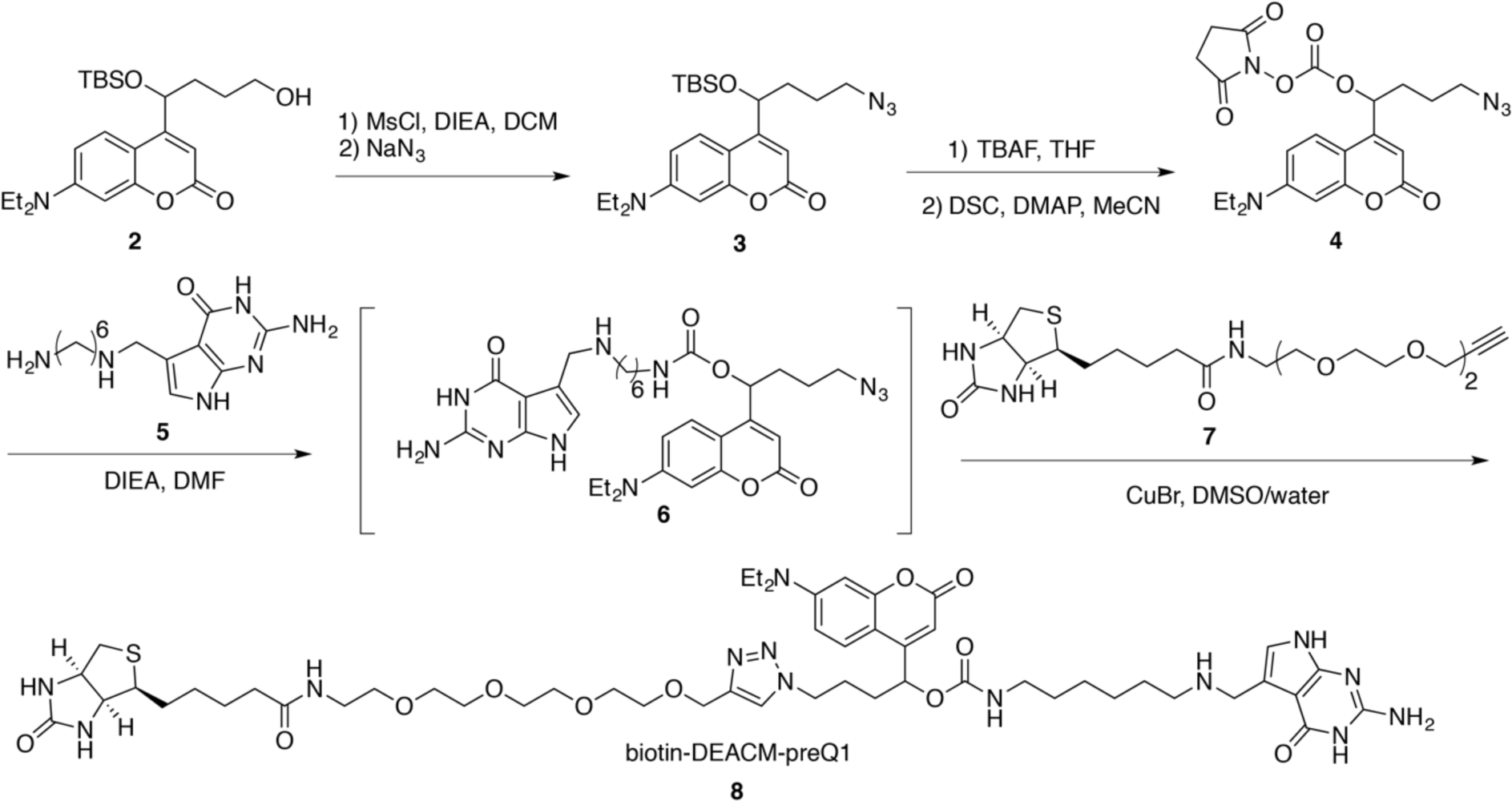
Synthesis of the preQ_1_-DEACM-biotin (8) Boc = *tert*-butyloxycarbonyl protecting group, MsCl = methanesulfonyl chloride, DCM = dichloromethane, TBS = tert-butyldimethylsilyl protective group, TBAF = tetra-*n*-butylammonium fluoride, THF = tetrahydrofuran, DSC = N-succinimidyl carbonate, DMAP = 4-dimethylaminopyridine, DIEA = N,N-diisopropylethylamine, DMF = dimethylformamide, DMSO = dimethyl sulfoxide.

Previously, we chose the photo-cleavable linker 6-bromo-7-aminoethoxycoumarin-4-ylmethoxycarbonyl (Bac) to synthesize the ‘photo-cage’, biotin-Bac-preQ_1_ **(1)** (Figure 1B). [32] To allow for sequential photo-activation of mRNA translation, an additional photo-sensitive linker that is responsive to a longer wavelength of light was desired. Inspired by previously reported work, [39] we chose to explore [7-(diethylamino)coumarin-4-yl]-methyl (DEACM) as an additional photo-activable linker that can be sequentially activated. The DEACM linker has a wide absorbance spectrum and was previously reported to be cleaved in cellular conditions by irradiation with 470 nm light. [40] Thus, the DEACM linker should form a sequentially photo-activable linker pair with our previously reported Bac linker, which is uncaged by irradiation with 365-405 nm light. To synthesize the new ‘photo-cage’ biotin-DEACM-preQ_1_ **(8)** (Scheme 1), the DEACM-based building block **(2)** was subjected to sequential mesylation and azidation to generate azide compound **(3)**. Next, the DEACM NHS-ester **(4)** was obtained through deprotection of the silica protecting group and N-succinimidyl carbonate (DSC) treatment. Subsequently, the preQ_1_ derivative **(5)** was coupled to the DEACM NHS-ester **(4)** to yield the preQ_1_-DEACM conjugate **(6)**. To introduce a biotin affinity handle to facilitate purification of the labeled mRNAs, the preQ_1_-DEACM conjugate **(6)** was further coupled with commercially available biotin-PEG_4_-alkyne **(7)** via click chemistry to obtain the final TGT enzymatic substrate biotin-DEACM-preQ_1_ **(8)**.

To demonstrate that biotin-Bac-preQ_1_ **(1)** and biotin-DEACM-preQ_1_ **(8)** can be sequentially released from RNA upon photo-irradiation with two wavelengths of light, *in-vitro* uncaging of labeled RNA oligos was performed using either 365 nm or 456 nm light followed by denaturing polyacrylamide gel electrophoresis (denaturing-PAGE) analysis. The ‘Tag’ oligo was used as RNA substrate for TGT labeling (Figure 2A). To covalently conjugate the photo-cage onto the ‘Tag’ oligo, TGT labeling using either biotin-Bac-preQ_1_ or biotin-DEACM-preQ_1_ as small molecule substrate was carried out in a dark room with minimal red ambient light to prevent undesired degradation of the light-sensitive preQ_1_ derivatives. Labeled ‘Tag’ oligo was further purified by streptavidin-biotin pull-down followed by denaturing-PAGE analysis. Covalent conjugation of the ‘photo-cage’ increased the molecular weight of the ‘Tag’ oligo, resulting in significant RNA band shifts shown in denaturing-PAGE (Figure 2B). [32-33] To trigger the cleavage of the ‘photo-cage’, the ‘Tag’ oligo labeled with biotin-DEACM-preQ_1_ was irradiated with a 456 nm lamp, while the ‘Tag’ oligo labeled with biotin-Bac-preQ_1_ was irradiated with a 365 nm lamp. We observed that upon irradiation with 456 nm light, only biotin-DEACM-preQ_1_ was released from the ‘Tag’ oligo, not the biotin-Bac-preQ_1_. Biotin-Bac-preQ_1_ was released from the ‘Tag’ oligo upon irradiation with 365 nm light. Thus, by using 456 nm and 365 nm light sources, sequential release of the ‘photo-cages’ from RNA was achieved.

**Figure 2.**
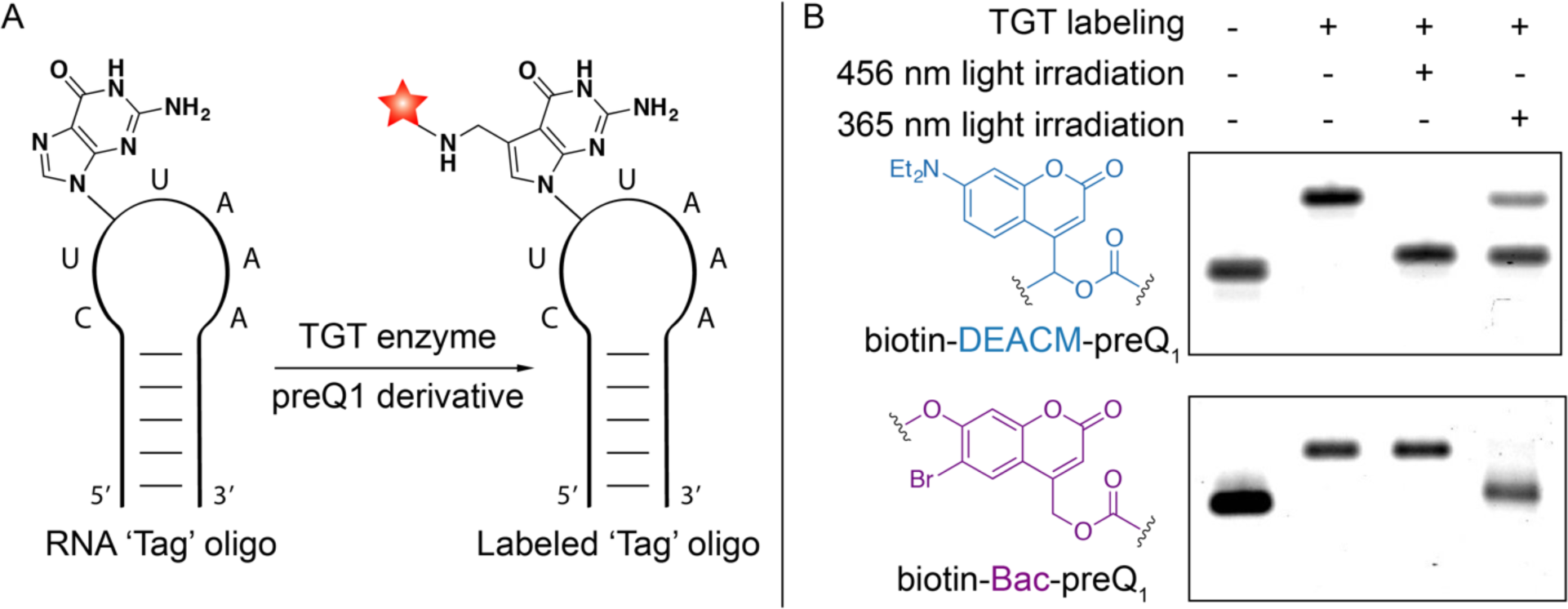
TGT labeling of the RNA ‘Tag’ oligo and *in-vitro* sequential photo-uncaging. A) *In-vitro* conjugation of RNA ‘Tag’ oligo with preQ_1_ derivatives through RNA-TAG. B) Denaturing 18% PAGE analysis of enzymatic labeling and subsequent photo-uncaging of the RNA ‘Tag’ oligos. Compared to the unlabeled RNA oligo, an upper gel shift in the second column demonstrates covalent attachment of the corresponding ‘photo-cage’ onto the RNA oligo. Lower gel shift in the third or fourth column demonstrates photo-release of the ‘photo-cage’ from the RNA oligo.

Having demonstrated that biotin-DEACM-preQ_1_ and biotin-Bac-preQ_1_ can be sequentially released from RNA oligos *in-vitro*, we examined whether these ‘photo-cages’ can be sequentially photo-released from mRNA in live mammalian cells. Mature IVT-mRNAs coding for either the green fluorescent protein (GFP) or the red fluorescent protein (RFP), with three TGT labeling sites located along the 5′-UTR, were synthesized following a previously reported mRNA transcription and maturation protocol. [32] Using TGT enzymatic labeling, the mature IVT-mRNA coding for GFP was conjugated with biotin-DEACM-preQ_1_ while the IVT-mRNA coding for RFP was conjugated with biotin-Bac-preQ_1_. To get rid of unlabeled IVT-mRNAs, biotin-streptavidin affinity purification was performed (60% recovery). Equal amounts of photocaged IVT-mRNAs coding for GFP or RFP were co-transfected into HEK-293 cells using lipofectamine reagent (Figure 3A). mRNA translation activity was quantified by fluorescence imaging. Minimal translation activity was observed for the photocaged mRNAs, shown as dark cells (Figure 3B). Two hours post transfection, selected cells were irradiated with either 488 nm or 405 nm wavelength of laser light to trigger the release of the ‘photo-cages’ from mRNA. To allow time for sufficient mRNA translation and maturation, cells were imaged 8 hours after laser irradiation. As expected from our *in-vitro* studies, the 488 nm irradiation only triggered the release of the biotin-DEACM-preQ_1_ from mRNA. As a result, recovered protein expression of GFP was observed only in cells that were irradiated with the 488 nm laser (Figure 3B). Importantly, expression of RFP was not observed in these cells, which was expected because the biotin-Bac-preQ_1_ used to cage the RFP-mRNA is not responsive to 488 nm irradiation. In contrast, expression of both GFP and RFP was observed in cells that were irradiated with 405 nm wavelength of laser, because both biotin-Bac-preQ_1_ and biotin-DEACM-preQ_1_ have absorbance at 405 nm wavelength. These observations were consistent with our *in-vitro* uncaging experiments, that the longer wavelength of light only triggered the release of biotin-DEACM-preQ_1_, whereas the shorter wavelength of light triggered the release of both ‘photocages’ from mRNA. Moreover, expression of fluorescent protein was only observed in laser irradiated cells, not in adjacent cells, demonstrating photo-activation of gene expression with high cellular resolution.

**Figure 3.**
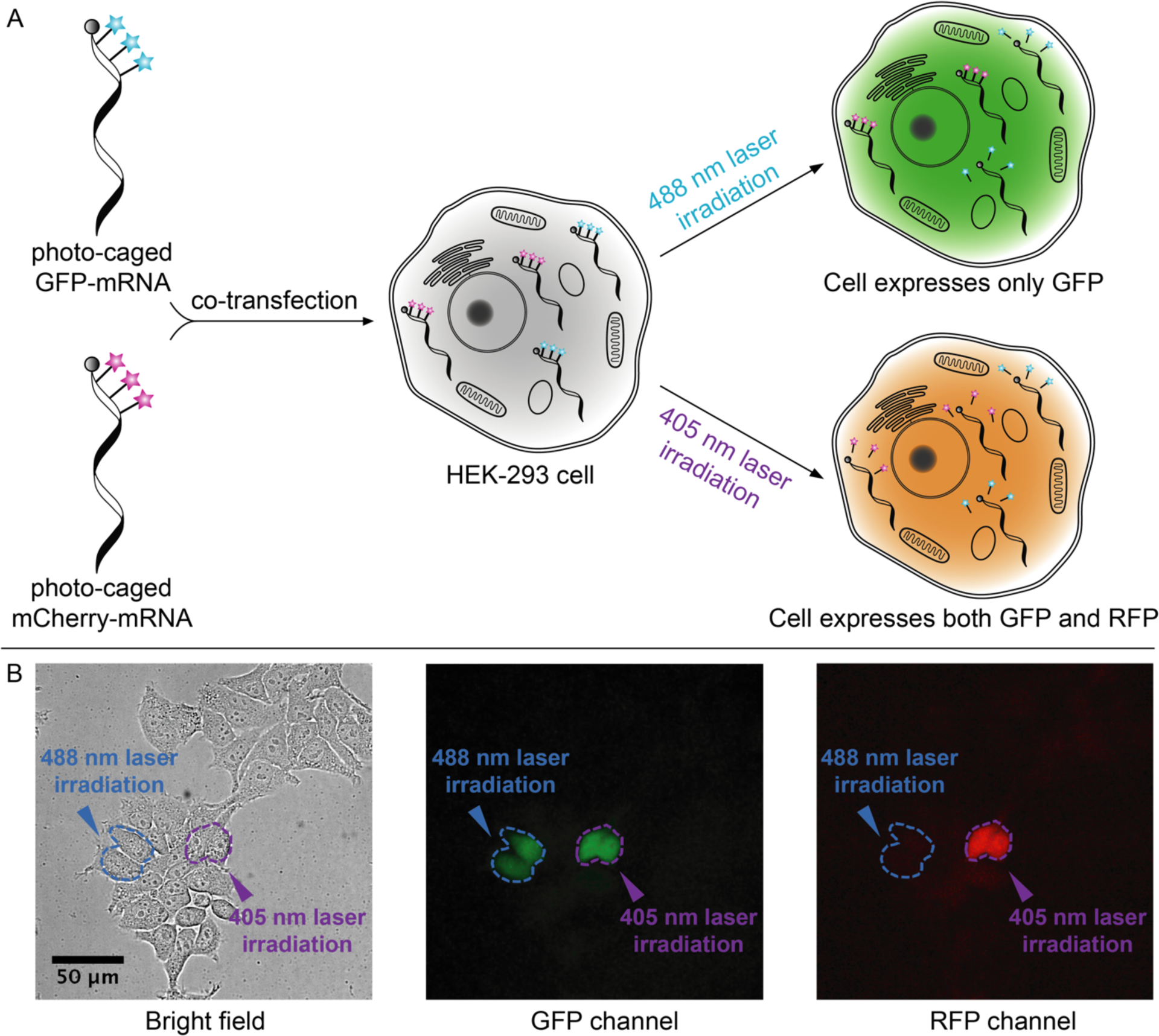
Live cell photo-activation of mRNA translation. A) HEK-293 cells are co-transfected with caged GFP-mRNA and caged mCherry-mRNA, followed by photo-uncaging with either 488 nm or 405 nm laser irradiation. B) Live cell fluorescence imaging. Selected cells (circled in blue) irradiated with 488 nm laser only express GFP, shown as green cells in the GFP channel and dark cells in the RFP channel. Selected cells (circled in purple) irradiated with 405 nm light express both GFP and RFP. Scale bar = 50 µm.

Next, we demonstrated sequential photo-activation of two mRNAs in the same living cell. Caged IVT-mRNAs coding for GFP and RFP were co-transfected into HEK-293 cells. Two hours post transfection, the selected cell was first irradiated with 488 nm laser light to trigger the release of biotin-DEACM-preQ_1_ from GFP-mRNA. Five hours post transfection, the same cell was then irradiated with 405 nm laser light to trigger the release of the biotin-Bac-preQ_1_ from RFP-mRNA. Cells were continuously imaged to quantify protein expression level (Figure 4A). As shown in the fluorescence images, the first irradiation with 488 nm laser light triggered the expression of GFP, while the expression of RFP remained silenced. As expected, the subsequent irradiation with 405 nm laser light activated the expression of RFP. GFP and RFP expression levels in laser irradiated cells were quantified by measuring fluorescence intensity (Figure 4B). Using this model, we demonstrated sequential photo-activation of two mRNAs in the same cell by using two wavelengths of lights.

**Figure 4.**
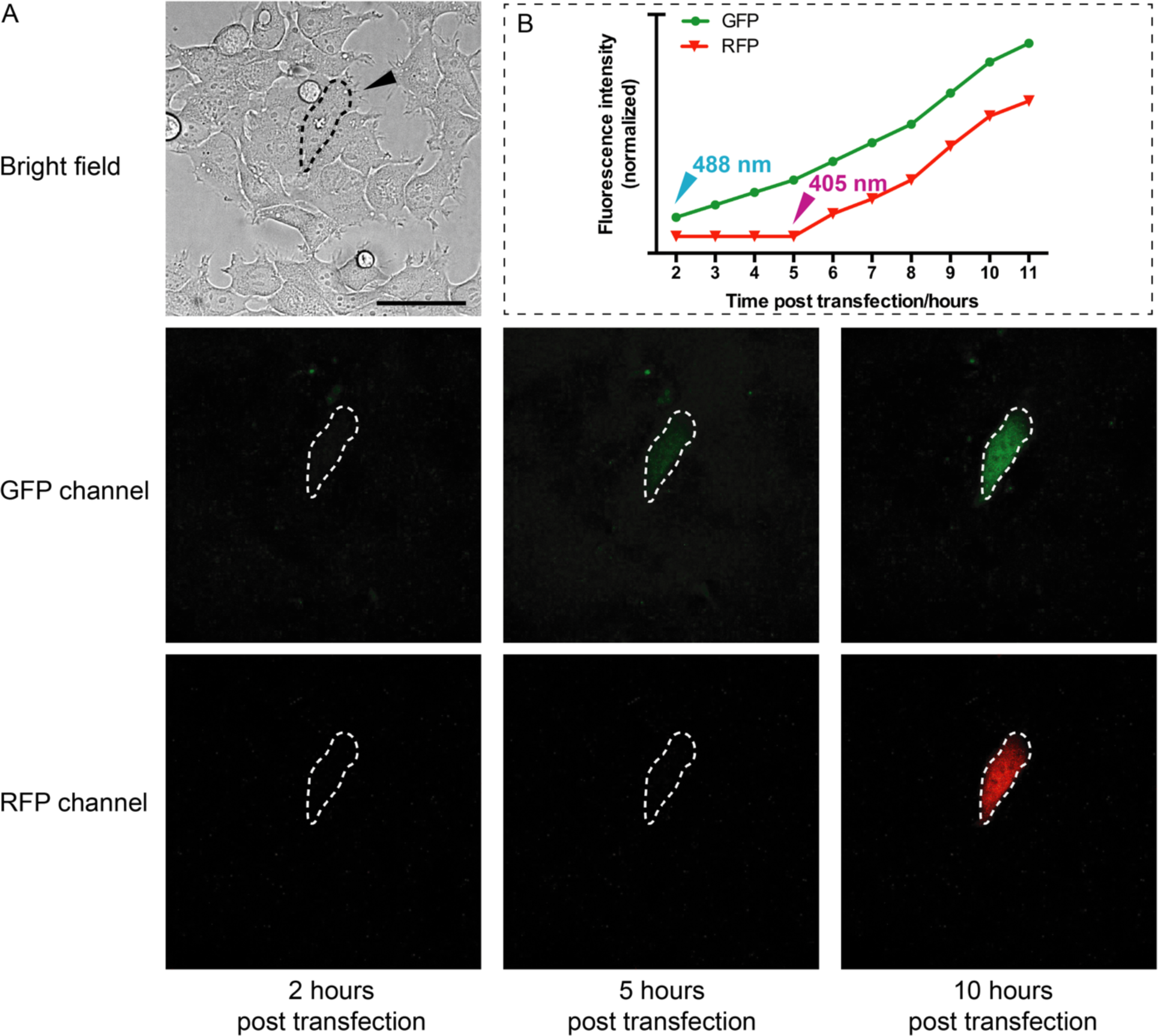
Live cell sequential photo-activation of two mRNAs within the same cell. A) A selected cell (circled in dash line) was first irradiated with 488 nm laser light to activate GFP-mRNA. Three hours after the first photo-uncaging event, the same cell was irradiated with 405 nm laser light to activate RFP-mRNA (Scale bar = 50 µm). B) Normalized average fluorescence intensity of GFP and RFP from this selected cell is plotted against time (hours post transfection).

## CONCLUSION

In conclusion, we have developed a technique that allows sequential photo-activation of two mRNAs with single-cell resolution. We demonstrated the synthesis and photochemical properties of two ‘photo-cages’, biotin-Bac-preQ_1_ **(1)** and biotin-DEACM-preQ_1_ **(8)**. These ‘photocages’ were covalently and site-specifically conjugated onto the 5′-UTR of mRNA through TGT enzymatic labeling. As a result, translation efficiency of the labeled mRNA was severely diminished compared to the unlabeled mRNA. This pair of ‘photo-cages’ can be released from mRNA transcripts sequentially upon irradiation with 365 nm/405 nm light (lamp excitation) or 405 nm/488 nm light (laser excitation), leading to translational activation of the corresponding mRNA. Irradiation with a longer wavelength of light only cleaves one ‘photo-cage’ (biotin-DEACM-preQ_1_) from RNA, while a shorter wavelength of light cleaves both ‘photo-cages’. By using the appropriate order of photo-irradiation, sequential photo-activation of two mRNAs within the same cell was demonstrated by live cell fluorescence imaging. We believe that the ability to sequentially photo-activate two genes with high spatial-temporal resolution provides a powerful and versatile optogenetic tool to build robust, complex, and scalable synthetic gene networks. Such a tool may improve capabilities to precisely manipulate biological networks, which can aid studies of gene regulatory mechanisms, promote the engineering of artificial biological systems, and facilitate the development of novel therapeutic applications.

## AUTHOR INFORMATION

### Notes

The authors declare no competing financial interest.

## ACKNOWLEDGMENTS

The project or effort depicted is sponsored by the Defense Advanced Research Projects Agency Biological Technologies Office (BTO) Safe Genes Program under Contract Number HR0011-18-2-0039. The content of the information does not necessarily reflect the position or the policy of the government, and no official endorsement should be inferred.

