## Supplementary material for "Multiplexed photo-activation of mRNA with single-cell resolution": SI.pdf

#### Abstract

We demonstrate sequential optical activation of two types of mRNAs in the same mammalian cell through the sequential photocleavage of small molecule caging groups (‘photo-cages’) tethered to the 5′ untranslated region (5′-UTR) of an mRNA. Synthetic ‘photo-cages’ were conjugated onto target mRNA using RNA-TAG, an enzymatic site-specific RNA modification technique. Translation of mRNA was severely reduced upon conjugation of the ‘photo-cages’ onto the 5′-UTR. However, subsequent photo-release of the ‘cages’ from the mRNA transcript triggered activation of translation with single-cell spatiotemporal resolution. To achieve sequential photo-activation of two mRNAs in the same cell, we synthesized a pair of ‘photo-cages’ which can be selectively cleaved upon photo-irradiation with different wavelengths of light. Sequential photo-activation of two mRNAs enabled precise optical control of translation of two unique transcripts. We believe that this modular approach to precisely and rapidly control gene expression will serve as a powerful tool in future biological studies that require controlling translation of multiple transcripts with high spatiotemporal resolution.

### General Materials

#### Reagents and instruments

Commercially available methanesulfonyl chloride, sodium azide, tetra-*n*-butylammonium fluoride in THF (1M), *N*-succinimidyl carbonate, 4-dimethylaminopyridine, *N*,*N*-diisopropylethylamine, copper(I) bromide and common organic solvents were obtained from Sigma-Aldrich. Deuterated chloroform ($CDCl_3$) was obtained from Cambridge Isotope Laboratories. All reagents obtained from commercial suppliers were used without further purification. Analytical thin-layer chromatography was performed on E. Merck silica gel 60 $F_{254}$ plates. Silica gel flash chromatography was performed using E. Merck silica gel (type 60SDS, 230-400 mesh). Solvent mixtures for chromatography are reported as v/v ratios. HPLC analysis was carried out on an Eclipse Plus C8 analytical column with *Phase A*/*Phase B* gradients [*Phase A*: $H_2O$ with 0.1% formic acid; *Phase B*: MeOH with 0.1% formic acid]. HPLC purification was carried out on Zorbax SB-C18 semipreparative column with *Phase A*/*Phase B* gradients [*Phase A*: $H_2O$ with 0.1% formic acid; *Phase B*: MeOH with 0.1% formic acid]. Proton nuclear magnetic resonance ($^1$H NMR) spectra were recorded on a VarianVX-500 MHz spectrometer, and were referenced relative to residual proton resonances in $CDCl_3$ (at 7.24 ppm). Chemical shifts were reported in parts per million (ppm, $\delta$) relative to tetramethylsilane (at 0.00 ppm). $^1$H NMR splitting patterns are assigned as singlet (s), doublet (d), triplet (t), quartet (q) or pentuplet (p). All first-order splitting patterns were designated on the basis of the appearance of the multiplet. Splitting patterns that could not be readily interpreted are designated as multiplet (m) or broad (br). Carbon nuclear magnetic resonance ($^{13}$C NMR) spectra were recorded on a Varian VX-500 MHz spectrometer, and were referenced relative to residual proton resonances in $CDCl_3$ (at 77.23 ppm). Electrospray Ionization-Time of Flight (ESI-TOF) spectra were obtained on an Agilent 6230 Accurate-Mass TOF mass spectrometer.

DNA oligonucleotides were purchased from Integrated DNA Technologies (Coralville, IA) and Eton Bioscience (California, CA). Molecular biology reagents such as restriction digestion enzymes, Q5 DNA polymerase, T7 RNA polymerase, Vaccinia Capping System, *E. coli* Poly(A) Polymerase, nucleotide stains and competent bacterial strains were purchased from New England Biolabs (Ipswitch, MA), Promega (Madison, WI), or Life Technologies (Carlsbad, CA). Dynabeads™ M-280 Streptavidin was purchased from Thermo Fisher Scientific (Waltham, MA). Fluorescence microscopy imaging was performed on an Axio Observer Z1 inverted microscope (Carl Zeiss Microscopy Gmb, Germany) with Yokogawa CSU-X1 spinning disk confocal unit using a 20 x, 0.8 NA objective to an ORCA-Flash4.0 V2 Digital CMOS camera (Hamamatsu, Japan). Fluorophores were excited with laser diodes (405 nm; 20 mW, 488 nm; 30 mW). Photo-uncaging was performed using a DirectFRAP module and the 405 nm and 488 nm laser (Carl Zeiss). Images were acquired using Zen Blue software (Carl Zeiss) and processed using Image J.

#### Reaction Buffers

TGT Storage Buffer: 25 mM HEPES, pH 7.3, 2 mM DTT, 1 mM EDTA, and 100 µM PMSF.
TGT Reaction Buffer: 100 mM HEPES, pH 7.3, 5 mM DTT, and 20 mM MgCl2.
T7 Reaction Buffer: 40 mM Tris pH 7.5, 5 mM DTT, 25 mM MgCl2, 2 mM spermidine.
Dynabeads™ streptavidin binding and washing (B&W) buffer 2X: 10 mM Tris-HCl (pH 7.5), 1mM EDTA, 2 M NaCl.

Dynabeads™ streptavidin binding and washing buffer with Tween (BW&T) 2X: 10 mM Tris-HCl (pH 7.5), 1mM EDTA, 2 M NaCl, 0.1% Tween 20.

Dynabeads™ streptavidin solution A: DEPC-treated 0.1 M NaOH, DEPC-treated 0.05 M NaCl.

Dynabeads™ streptavidin solution B: DEPC-treated 0.1 M NaCl.

Dynabeads™ streptavidin elution buffer: 950 μL formamide, 20 μL 500 mM EDTA, 30 μL RNase free water.

#### Chemical Synthesis

##### Synthesis of compound **(3)**

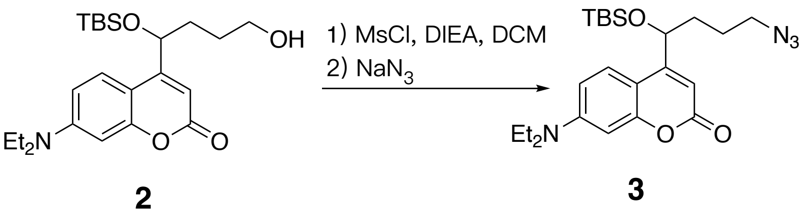

Scheme S1. Synthesis of compound **(3)**

4-(4-azido-1-((tert-butyldimethylsilyl)oxy)butyl)-7-(diethylamino)-2H-chromen-2-one **(3)**

A solution of the previously reported 4-(1-((*tert*-butyldimethylsilyl)oxy)-4-hydroxybutyl)-7-(diethylamino)-2*H*-chromen-2-one **(2)** (50.0 mg, 0.12 mmol) in DCM (5 mL) was treated with DIEA (29.8 mg, 0.24 mmol) and MsCl (20.6 mg, 0.18 mmol). [1, 2] Then the reaction mixture was stirred for 12 hours at room temperature. Afterwards, the reaction solution was quenched with water in ice bath and extracted with 5 mL DCM for 3 times. The combined organic solution was washed with brine and dried with $Na_2SO_4$. All organic solvent was removed *in vacuo*. The resulting product was dissolved in DMF (3 mL). Next, $NaN_3$ (15 mg, 0.24 mmol) was added to the solution. The reaction solution was heated at 80°C for 5 hours. After cooling the reaction to room temperature, water (10 mL) was added to quench the reaction, followed by EtOAc (5mL*6) extraction. Then the organic solution was dried with $Na_2SO_4$ and removed *in vacuo*. The resulting crude material was then purified by flash chromatography (0-10% EtOAc in hexanes) to afford compound **(3)** as a yellow solid (43.2 mg, 81%). $^1$H NMR (500 MHz, $CDCl_3$, δ): 7.45 (d, *J* = 9.0 Hz, 1H), 6.57 (d, *J* = 9.1 Hz, 1H), 6.52 (d, *J* = 2.5 Hz, 1H), 6.18 (s, 1H), 4.93-4.88 (m, 1H), 3.45-3.37 (m, 4H), 3.37-3.23 (m, 2H), 1.92-1.74 (m, 2H), 1.74-1.66 (m, 2H), 1.21 (t, *J* = 7.1 Hz, 6H), 0.92 (s, 9H), 0.09 (s, 3H), –0.03 (s, 3H). $^{13}$C NMR (126 MHz, $CDCl_3$, δ): 162.47, 158.10, 156.56, 150.27, 125.04, 108.35, 105.87, 105.87, 97.90, 70.50, 51.27, 44.68, 44.68, 35.17, 25.77, 25.77, 25.77, 24.69, 18.15, 12.44, 12.44, – 4.70, –5.15. HRMS (M+H$^+$) calcd for $[C_{23}H_{37}N_4O_3Si]^+$ 445.2629, found 445.2633.

##### Synthesis of compound **(4)**

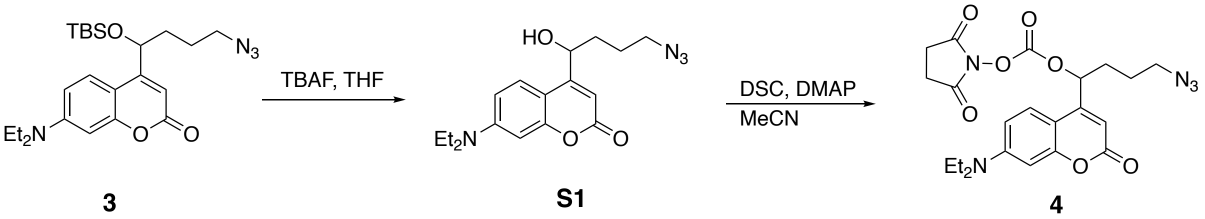

Scheme S2. Synthesis of compound **(4)**

4-(1-((*tert*-butyldimethylsilyl)oxy)-4-hydroxybutyl)-7-(diethylamino)-2*H*-chromen-2-one **(S1)**

A solution of compound **(3)** (20 mg, 0.045 mmol) in THF (5 mL) was treated with TBAF solution (1 M, 1.35 mmol). The resulting solution was stirred at room temperature for 6 hours. After the deprotection, the solution was directly subjected to semipreparative HPLC purification, using a C18 column [gradient of $H_2O$ with 0.1% formic acid and MeOH with 0.1% formic acid 95:5 (0 min) to 5:95 (10 min to 18min)]. Note: We tried to purify the compound by silica flash chromatography, which leads to complete decomposition of compound **(S1)**. The fractions containing product compound **(S1)** from semipreparative HPLC purification were dried *in vacuo* at room temperature on rotary evaporator. The resulting product was further dried *in vacuo* with mechanical pump for 1 hour at room temperature. Compound **(S1)** was obtained as yellow oil (18.1 mg, 96%). $^1$H NMR (500 MHz, $CDCl_3$, δ): 7.34 (d, *J* = 8.9 Hz, 1H), 6.54 (d, *J* = 8.8 Hz, 1H), 6.43 (s, 1H), 6.20 (s, 1H), 4.97-4.91 (m, 1H), 3.41 (s, 1H), 3.36-3.27 (m, 4H), 1.88 (m, 1H), 1.82-1.64 (m, 3H), 1.13 (t, *J* = 7.1 Hz, 6H). $^{13}$C NMR (126 MHz, $CDCl_3$, δ): 162.88, 158.64, 156.39, 150.26, 125.25, 109.17, 105.91, 105.26, 98.20, 77.35, 69.33, 51.24, 44.95, 33.84, 25.11, 12.51, 12.36. HRMS (M+Na+) calcd for [C17H22N4O3Na]+ 353.1384, found 353.1586.

4-azido-1-(7-(diethylamino)-2-oxo-2H-chromen-4-yl)butyl (2,5-dioxopyrrolidin-1-yl) carbonate **(4)**
Compound **(S1)** (18.1 mg, 0.043 mmol) was dissolved in MeCN, followed by addition of DSC (22.0 mg, 0.086 mmol) and DMAP (15.9 mg, 0.13 mmol). The reaction solution was stirred for 12 hours at room temperature, followed by semipreparative HPLC purification, using a C18 column [gradient of $H_2O$ with 0.1% formic acid and MeOH with 0.1% formic acid 95:5 (0 min) to 5:95 (10 min to 18 min)]. The fractions containing product **(4)** from semipreparative HPLC purification were dried *in vacuo* on rotary evaporator without heat. The resulting product was further dried *in vacuo* with mechanical pump for 1 hour at room temperature. Note: After dried in *in vacuo* at room temperature, there were small amount of unknown by-product generated. Thus, the NMR spectrums of compound **(4)** were not obtained. However, compound **(4)** is confirmed by low resolution mass spectrometry, using fresh fraction from semipreparative HPLC. LRMS ($M+H^+$) calcd for $[C_{22}H_{26}N_5O_7]^+$ 472.2, found 472.1.

##### Synthesis of compound **(8)**

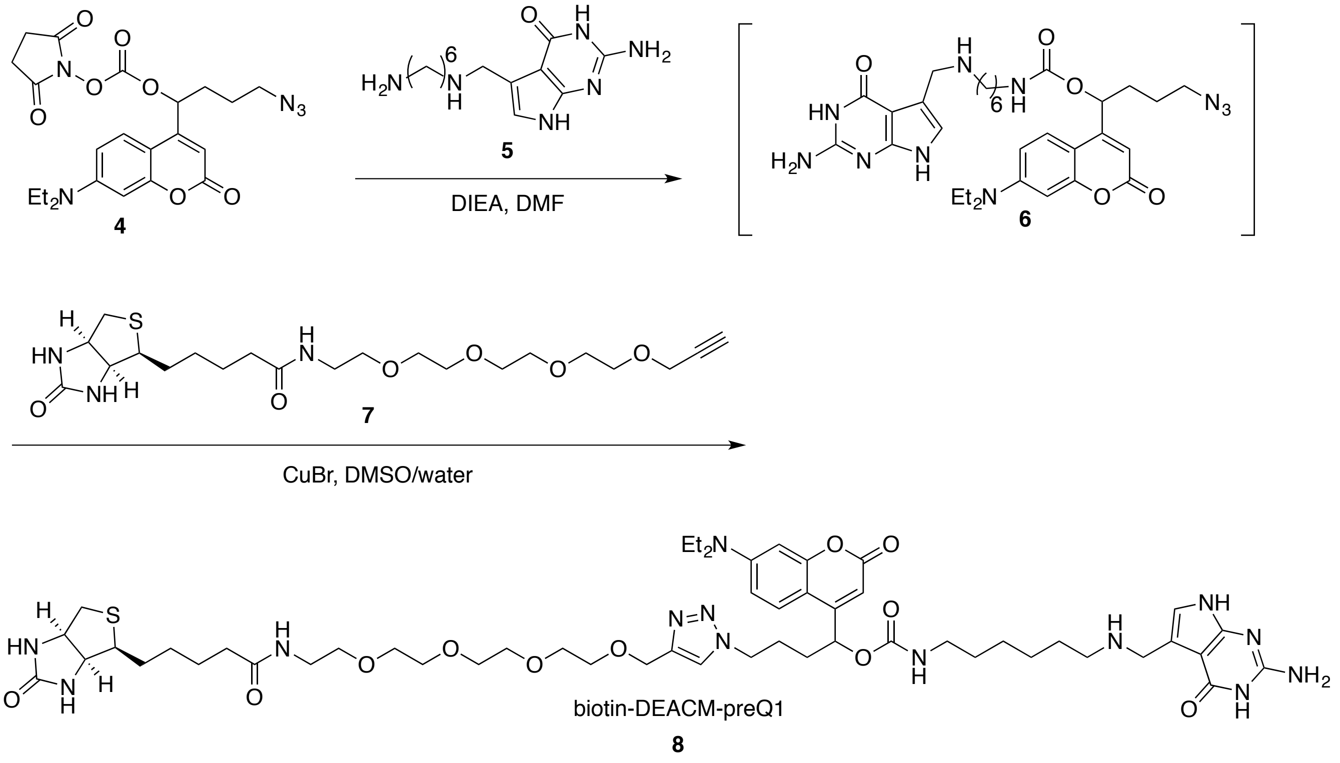

Scheme S3. Synthesis of compound **(8)**

1-(7-(diethylamino)-2-oxo-2*H*-chromen-4-yl)-4-(4-(15-oxo-19-((3a*S*,4*S*,6a*R*)-2-oxohexahydro-1H-thieno[3,4-d]imidazol-4-yl)-2,5,8,11-tetraoxa-14-azanonadecyl)-1*H*-1,2,3-triazol-1-yl)butyl (6-(((2-amino-4-oxo-4,7-dihydro-1*H*-pyrrolo[2,3-d]pyrimidin-5-yl)methyl)amino)hexyl)carbamate **(8)**

Compound **(4)** (5.0 mg, 0.011 mmol) was dissolved in DMF (0.5 mL) followed by slow addition of the previously reported compound **(5)** [3] (8.8 mg, 0.032 mmol in 0.5 mL DMF) and DIEA (7.1 mg, 0.055 mmol). The reaction solution was stirred for 1 hour at room temperature. The crude substitution product was directly subjected to semipreparative HPLC purification, using a C18 column [gradient of $H_2O$ with 0.1% formic acid and MeOH with 0.1% formic acid 95:5 (0 min) to 5:95 (10 min to 18min)]. Shielded from light, the semipreparative HPLC fractions containing product compound **(6)** (confirmed by low resolution mass spectrometry, LRMS ($M+H^+$) calcd for $[C_{31}H_{43}N_{10}O_5]^+$ 635.3, found 635.3) were dried *in vacuo* with ice bath. The resulting product was directly used for the next step of synthesis. Protected from light, CuBr (0.8 mg, 0.0055 mmol in DMSO:$H_2O$=0.3 ml:0.3 mL solution) was added to compound **(6)** followed by the addition of commercial available compound **(7)** (5.0 mg, 0.011 mmol in 0.4 mL DMSO; commercially available at BroadPharm, San Diego, USA; CAS# 1458576-00-5). The reaction mixture was stirred for 2 hours at room temperature, followed by semipreparative HPLC purification, using a C18 column [gradient of $H_2O$ with 0.1% formic acid and MeOH with 0.1% formic acid 95:5 (0 min) to 5:95 (10 min to 18min)]. Protected from light, the fractions containing compound **(8)** from semipreparative HPLC were dried *in vacuo* at room temperature. After further vacuum drying in ice bath, there was a trace amount of water left in the vial due to the low temperature. To completely dry the final product compound **(8)**, a solvent mixture of $H_2O$:MeCN=0.3 mL:0.3 mL was added to the vial containing compound **(8)**, followed by lyophilization. Compound **(8)** was obtained as white residue (0.6 mg, 5% for 2 steps). $^1$H NMR (500 MHz, $CDCl_3$, δ): 8.31-8.11 (m, 1H), 7.69-7.55 (m, 1H), 7.33-7.28 (m, 1H), 6.80-6.65 (m, 1H), 6.59-6.44 (m, 2H), 6.44-6.30 (m, 2H), 5.41-5.24 (m, 4H), 4.62-4.38 (m, 2H), 4.36-4.16 (m, 2H), 4.15-4.12 (m, 1H), 3.67-3.44 (m, 14H), 3.36-3.25 (4H), 3.21-2.98 (m, 2H), 2.91-2.67 (2H), 2.21-2.11 (m, 2H), 1.97-1.91 (m, 2H), 1.32-1.15 (m, 14H), 1.14-1.08 (m, 2H), 0.87-0.72 (m, 6H). $^{13}$C NMR (126 MHz, $CDCl_3$, δ): 178.98, 177.57, 174.73, 167.36, 165.51, 158.43, 154.33, 154.06, 150.28, 147.65, 131.75, 130.34, 130.06, 121.19, 120.50, 119.05, 109.78, 100.43, 98.61, 93.69, 84.55, 70.59, 70.55, 70.54, 70.53, 70.50, 70.43, 70.29, 70.25, 61.89, 61.77, 60.34, 51.40, 50.19, 45.07, 39.79, 39.46, 37.30, 36.23, 32.23, 29.98, 29.85, 29.64, 29.43, 27.53, 27.48, 25.82, 25.66, 23.01, 14.46, 12.86, 12.76. HRMS ($M+H^+$) calcd for $[C_{52}H_{78}N_{13}O_{11}S]^+$ 1092.5659, found 1092.5664.

#### Transcription template

##### GFP-multiTag transcription template

The GFP expression vector pcDNA3-GFP was purchased from Addgene (plasmid #13031) and as a gift from Doug Golenbock. Ultramer DNA oligo containing three active tags was purchased from Integrated DNA Technologies (San Diego, CA). GFP-multiTag vector was obtained by inserting the ultramer DNA oligo into 5′-UTR of the GFP coding region through Gibson DNA assembly technique (NEB, Ipswitch, MA). DH5a competent cells (Life Technologies, Carlsbad, CA) were transformed with the ligation product and screened against ampicillin on agar plates overnight. Colonies were selected and overgrown for 16 hours. The overgrowth was subjected to DNA extraction with a QIAGEN Plasmid Maxi Kit (QIAGEN, Venlo, Limburg Netherlands). Sequencing was performed to verify the inserted sequence.

GFP-multiTag sequence (from T7 promoter to 3’ XbaI restriction enzyme cut site) is shown below. The ‘Tag’ sequences are underlined.

TAATACGACTCACTATAGGG***GCAGACTGTAAATCTGC***AGACCCAAGCTTGGTAGGTCAGTTGCAGTTACCG
AGCTCGGATCCACTAGTAACGGCCGC***GCAGACTGTAAATCTGC***CAGTGTGCTAGTCAGACAGATGGAATTC
TGCAGATATCCATCACACTGGCGGCCGCTCGA***GCAGACTGTAAATCTGC***GATGGTGAGCAAGGGCGAGGA
GCTGTTCACCGGGGTGGTGCCCATCCTGGTCGAGCTGGACGGCGACGTAAACGGCCACAAGTTCAGCGTG
TCCGGCGAGGGCGAGGGCGATGCCACCTACGGCAAGCTGACCCTGAAGTTCATCTGCACCACCGGCAAGC
TGCCCGTGCCCTGGCCCACCCTCGTGACCACCCTGACCTACGGCGTGCAGTGCTTCAGCCGCTACCCCGA
CCACATGAAGCAGCACGACTTCTTCAAGTCCGCCATGCCCGAAGGCTACGTCCAGGAGCGCACCATCTTCT
TCAAGGACGACGGCAACTACAAGACCCGCGCCGAGGTGAAGTTCGAGGGCGACACCCTGGTGAACCGCAT
CGAGCTGAAGGGCATCGACTTCAAGGAGGACGGCAACATCCTGGGGCACAAGCTGGAGTACAACTACAAC
AGCCACAACGTCTATATCATGGCCGACAAGCAGAAGAACGGCATCAAGGTGAACTTCAAGATCCGCCACAA
CATCGAGGACGGCAGCGTGCAGCTCGCCGACCACTACCAGCAGAACACCCCCATCGGCGACGGCCCCGTG
CTGCTGCCCGACAACCACTACCTGAGCACCCAGTCCGCCCTGAGCAAAGACCCCAACGAGAAGCGCGATC
ACATGGTCCTGCTGGAGTTCGTGACCGCCGCCGGGATCACTCTCGGCATGGACGAGCTGTACAAGTAATCT
AGA

##### RFP-multiTag transcription template

Similarly, RFP-multiTag vector was prepared. RFP-multiTag mRNA vector sequence (from T7 promoter to 3’ restriction enzyme cut site) is shown below. The ‘Tag’ sequences are underlined.

TAATACGACTCACTATAGGG***GCAGACTGTAAATCTGC***AGACCCAAGCTTGGTAGGTCAGTTGCAGTTACCG
AGCTCGGATCCACTAGTAACGGCCGC***GCAGACTGTAAATCTGC***CAGTGTGCTAGTCAGACAGATGGAATTC
TGCAGATATCCATCACACTGGCGGCCGCTCGA***GCAGACTGTAAATCTGC***GATGGTGAGCAAGGGCGAGGA
GGATAACATGGCCATCATCAAGGAGTTCATGCGCTTCAAGGTGCACATGGAGGGCTCCGTGAACGGCCACG
AGTTCGAGATCGAGGGCGAGGGCGAGGGCCGCCCCTACGAGGGCACCCAGACCGCCAAGCTGAAGGTGA
CCAAGGGTGGCCCCCTGCCCTTCGCCTGGGACATCCTGTCCCCTCAGTTCATGTACGGCTCCAAGGCCTAC
GTGAAGCACCCCGCCGACATCCCCGACTACTTGAAGCTGTCCTTCCCCGAGGGCTTCAAGTGGGAGCGCG
TGATGAACTTCGAGGACGGCGGCGTGGTGACCGTGACCCAGGACTCCTCCCTGCAAGACGGCGAGTTCAT
CTACAAGGTGAAGCTGCGCGGCACCAACTTCCCCTCCGACGGCCCCGTAATGCAGAAGAAGACGATGGGC
TGGGAGGCCTCCTCCGAGCGGATGTACCCCGAGGACGGCGCCCTGAAGGGCGAGATCAAGCAGAGGCTG
AAGCTGAAGGACGGCGGCCACTACGACGCTGAGGTCAAGACCACCTACAAGGCCAAGAAGCCCGTGCAGC
TGCCCGGCGCCTACAACGTGAACATCAAGTTGGACATCACCTCCCACAACGAGGACTACACCATCGTGGAA
CAGTACGAACGCGCCGAGGGCCGCCACTCCACCGGCGGCATGGACGAGCTGTACAAGTAATATCTAACAC
AGACTCTCGGTACCATCATTTTCATATCCCCACCACCATCATTTAATGAATTCCATCAGGAATCCCTCACTTAA
AGCCCGCCGAAAGGCGGGCTTTTCTGTGTCTGCAGACTGGCCGTCGTTTTACACTCGAGCATGCATCTAGA

#### *In vitro* mRNA transcripts synthesis

##### *In vitro* transcription (IVT) reaction

GFP-multiTag vector was linearized using XbaI restriction digestion enzyme (NEB, Ipswitch, MA) to ensure uniform length transcription. For each IVT reaction, 10 µg of DNA plasmid was dissolved in cutsmart buffer (NEB,

Ipswitch, MA) to a final concentration of 300 ng/μL. 40 units of XbaI restriction digestion enzyme was added into the reaction mixture. The reaction was carried out at 37 °C for 4 hours to ensure complete digestion. The reaction solution was allowed to cool down to room temperature. The DNA product was extracted with an equal volume of molecular biology grade phenol / chloroform / isoamyl alcohol (25:24:1) and vortexed for 2 minutes followed by a 5-minute centrifugation at 10,000 RCF. The top aqueous layer was transferred to a fresh tube and an equal volume of chloroform was added. The mixture was vortexed for 2 minutes followed by a 5 minute centrifugation at 10,000 RCF. The aqueous layer was transferred into a fresh tube for DNA precipitation. The linearized DNA was precipitated by the addition of 0.1x volumes of 3M sodium acetate (pH = 5.2) and 2.5x volumes of 95% ethanol. The sample was chilled to -20 °C overnight, centrifuged at 16,100 RCF for 20 minutes at 4 °C, followed by gently removing ethanol. The DNA pellet was air dried and resuspended in 200 μL of RNase free water. The obtained DNA solution was used directly for IVT reaction. Each IVT reaction was set up with 50 ng/μL of linearized DNA template, 5 mM of each NTPs (ATP, CTP, UTP), 9 mM of GTP (NEB, Ipswitch, MA), 0.004 unit/μL of thermostable inorganic pyrophosphatase (NEB, Ipswitch, MA), 0.25 μg/μL T7 RNA polymerase, 0.05% Triton X-100 (Sigma, St. Louis, MO) and 1 unit/μL RNase Inhibitors, Murine (NEB, Ipswitch, MA). The IVT reaction was carried out at 37 °C for 4 hours to allow for sufficient RNA synthesis. To remove DNA template, 2 μL of 100 mM $CaCl_2$ and 20 units of Turbo DNase (Life Technologies, Carlsbad, CA) were added to the mixture and incubated at 37 °C for 1 hour. The mixture was then centrifuged at 10,000 RCF for 5 minutes at room temperature to pellet any remaining magnesium pyrophosphate. The supernatant was resuspended in 200 μL RNase free water. To precipitate the *in vitro* transcribed mRNA (IVT-mRNA) product, the solution was added with 100 μL of 8 M LiCl and chilled to -20 °C for 4 hours, followed by 20 minutes of centrifugation at 16,000 RCF at 4 °C. The supernatant was removed gently. The remaining RNA pellet was resuspended in 200 μL RNase free water and quantified at 260 nm. The IVT-mRNA was confirmed as a single observable UV shadowing band by 4% denaturing PAGE (4% polyacrylamide in TBE with 8M urea) and kept frozen at -20 °C until used.

##### Maturation of *in vitro* transcribed mRNA (IVT-mRNA)

Vaccinia capping system (NEB, Ipswitch, MA) was used to add a 7-methylguanylate cap structure (Cap 0) to the 5′ end of IVT-mRNA to allow 5′-cap dependent translation *initiation.* IVT-mRNA was diluted in 29.2 μL RNase free water with a final concentration of 0.5 μg/μL. The RNA solution was heated to 65 °C and held for 5 minutes to denature RNA. The mixture was then chilled on ice and held for another 5 minutes, followed by the addition of 4 μL of 10x capping buffer, 2 μL of 10 mM GTP, 2 μL of 4 mM S-adenosylmethionine (SAM), 32 units of RNase inhibitor and 2 μL of Vaccinia Capping Enzyme. The reaction mixture was incubated at 37 °C for 45 minutes. The capped IVT-mRNA product was purified by LiCl precipitation and directly used for polyadenylation. *E. coli* Poly(A) Polymerase (NEB, Ipswitch, MA) was used to polyadenylate IVT-mRNA. 30 μL of reaction mixture containing 15 μg of capped IVT-mRNA, 1 mM ATP, 30 units of RNase inhibitor was prepared in 1x *E. coli* Poly(A) polymerase reaction buffer and incubated at 37 °C for 1 hour. The maturated IVT-mRNA was purified by LiCl precipitation and directly used for *in vitro* TGT labeling.

#### TGT *in vitro* labeling and purification

##### TGT labeling of IVT mRNA transcript

TGT labeling conditions were adapted from our previous study (Figure S1). [3] Synthesized TGT substrate compound **7** was dissolved in RNase free water to obtain a concentration of 650 μM. For *in vitro* TGT reaction, 1 μM of IVT-mRNA, 50 μM of compound **7**, 2 unit/μL RNase inhibitor and 1.5 μM of TGT enzyme was assembled in 1x TGT reaction buffer. The reaction mixture was incubated at 37 °C for 4 hours. The crude labeled RNA transcript was purified by LiCl precipitation.

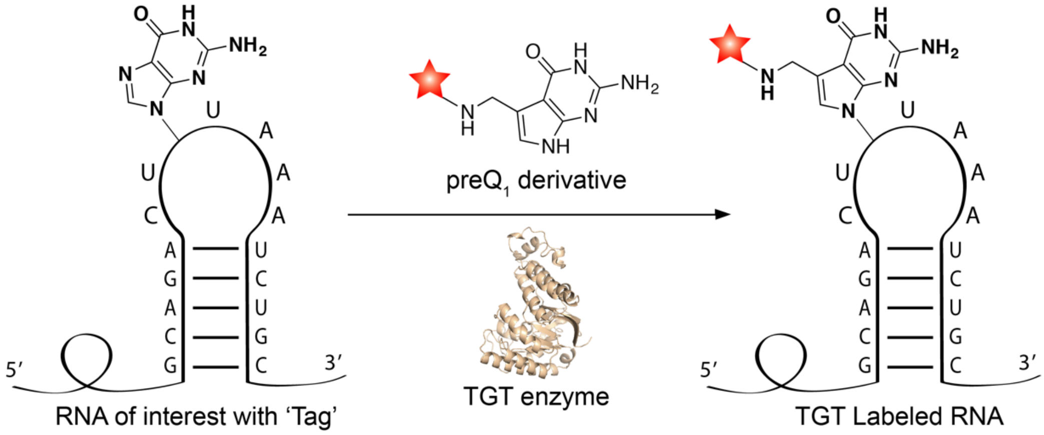

Figure S1. RNA labeling through 'RNA-TAG'. The bacterial tRNA guanine transglycosylase (TGT) exchanges a guanine nucleobase within the 17-nucleotide RNA stem-loop structure ('Tag') with a synthetic substrate ($preQ_1$ derivative) carrying a small-molecule functioning group (shown as red star).

##### Purification of TGT labeled mRNA transcript

The crude TGT labeling product was further purified using Dynabeads™ M-280 Streptavidin. Dynabeads™ M-280 Streptavidin beads stock solution (Thermo Fisher Scientific, Waltham, MA) was vortexed for 30 seconds. 150 μL of the well-mixed stock solution was transferred to a 2 mL Eppendorf tube. Following manufacture's protocal, the magnetic beads were washed with 300 μL of 1x BW&T buffer for 3 times, 300 μL of solution A for 2 times, 300 μL of solution B for 2 times and 300 μL of 1x BW&T buffer for 2 times After washing, the beads were incubated with labeled IVT-mRNA (use amount if available) in 1x BW&T buffer for 25 minutes to allow for the binding of IVT-mRNA to the magnetic beads. The solution was then rotated at room temperature for 25 minutes to allow for the binding of IVT-mRNA and the magnetic beads. The RNA bound beads were washed with 300 μL of 1x BW&T buffer 2 times and 300 μL of 1x B&W buffer 1 time. The beads were then resuspended in 100 μL elution buffer and incubated at 65 °C for 3 minutes, followed by centrifugation at 7,000 RCF for 1 minute. The resulting supernatant was then subjected to ethanol precipitation with the addition of 0.1x volumes of 3M sodium acetate and 4x volumes of ethanol. The purified labeled IVT-mRNA was stored in - 20 °C until used.

#### Live cell imaging and photo-uncaging

##### IVT mRNA transfection

HeLa cells (ATCC, Manassas, VA) were cultured in DMEM media (Life Technologies, Carlsbad, CA) with 10% FBS and P/. HeLa cells were plated at an initial density of around 40,000 cells per well in a Nunc Lab-Tek 8 well chamber slide (Thermo Scientific, Waltham, MA). Cells were allowed to adhere overnight, washed with Opti-MEM media (Life Technologies, Carlsbad, CA) and subsequently transfected in Opti-MEM media with an addition of 400 ng of IVT-mRNA and 1.0 μL of Lipofectamine® RNAiMAX (Life Technologies, Carlsbad, CA). Cells were transfected for 4 hours before washed with DMEM to remove any transfection reagents. Cells were allowed to grow in DMEM at 37 °C overnight to allow for sufficient protein expression.

##### Fluorescence microscopy imaging

All images, unless otherwise indicated, were acquired on a Yokagawa spinning disk system (Yokagawa, Japan) built around an Axio Observer Z1 motorized inverted microscope (Carl Zeiss Microscopy GmbH, Germany) with a 20x 1.42 NA objective to an Evolve 512x512 EMCCD camera (Photometrics, Canada) using ZEN imaging software (Carl Zeiss Microscopy GmbH, Germany). GFP was excited with a 488 nm, 100 mW OPSL laser, green. GFP expression from the IVT-mRNA was quantified by measuring average fluorescence level among more than 80 HeLa cells. Cells images were processed using ImageJ software.

#### NMR Spectrums

##### Compound **(3)**

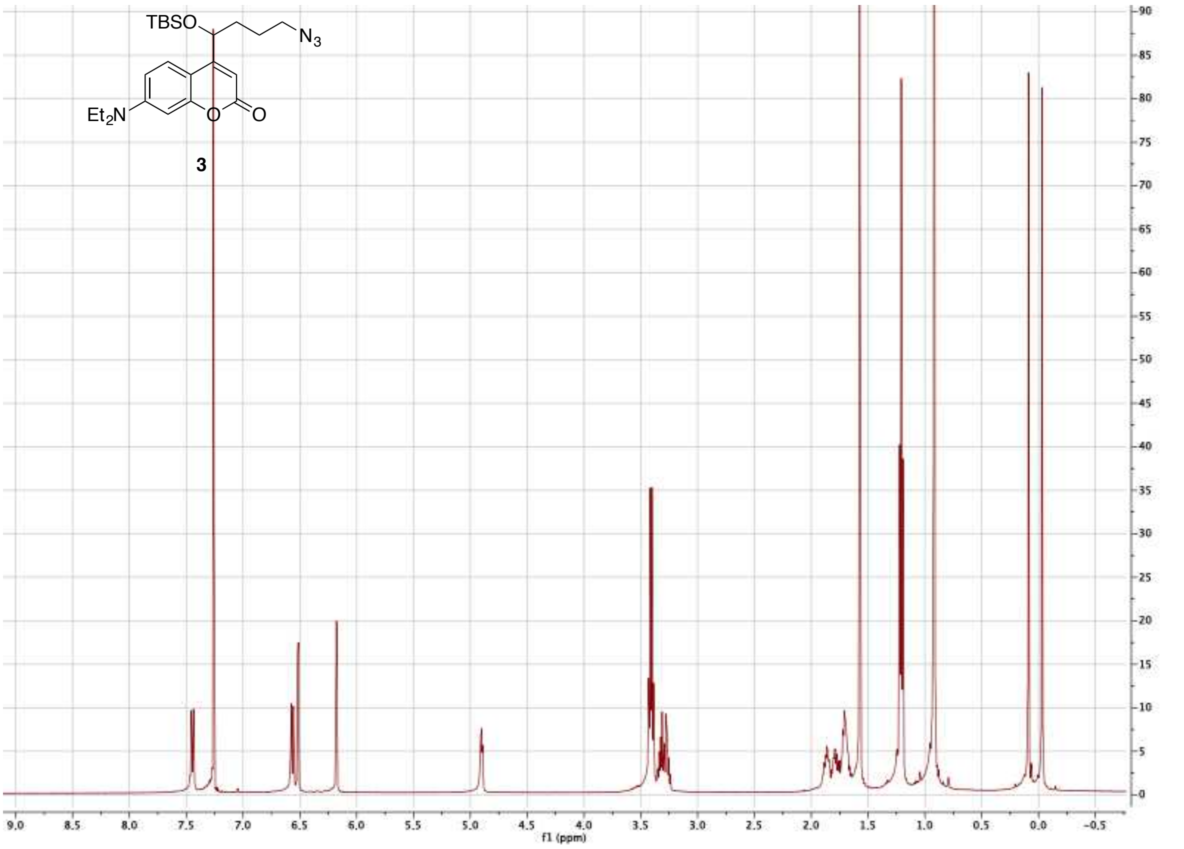

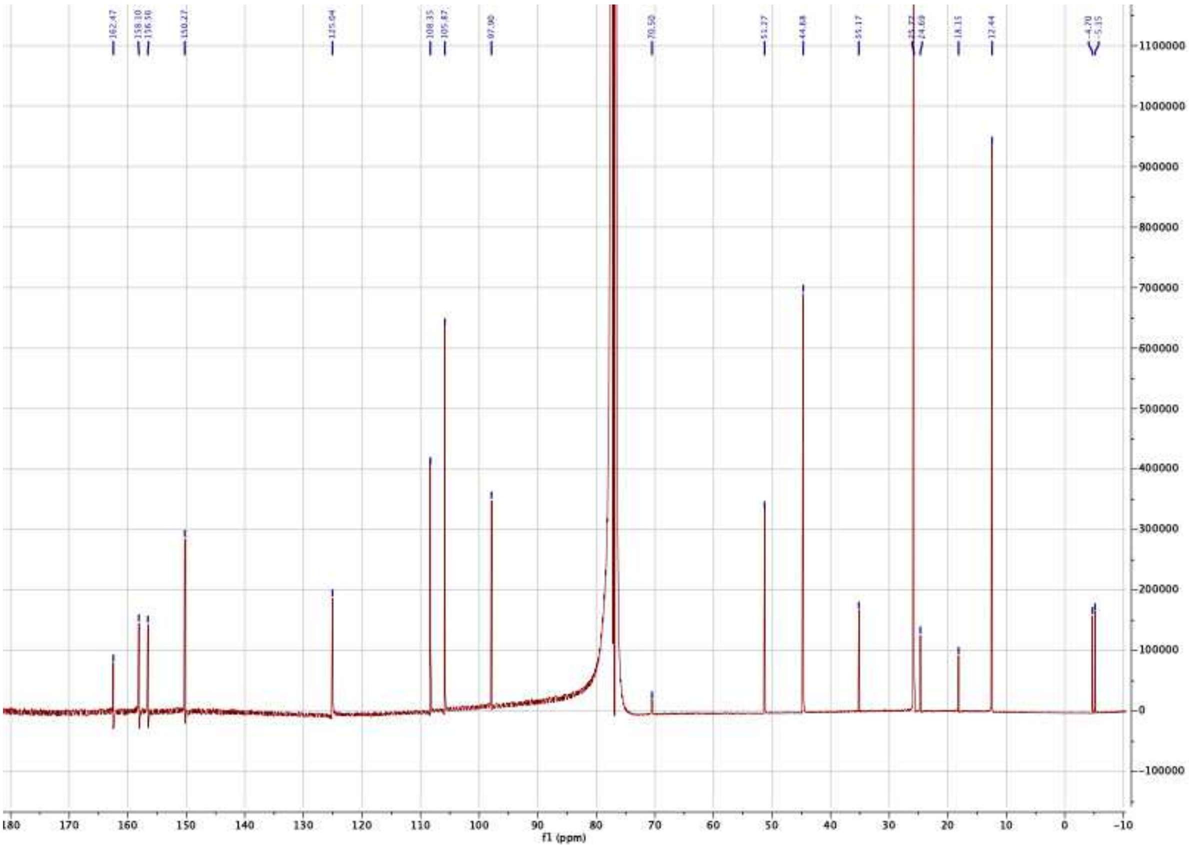

#### Compound **(S1)**

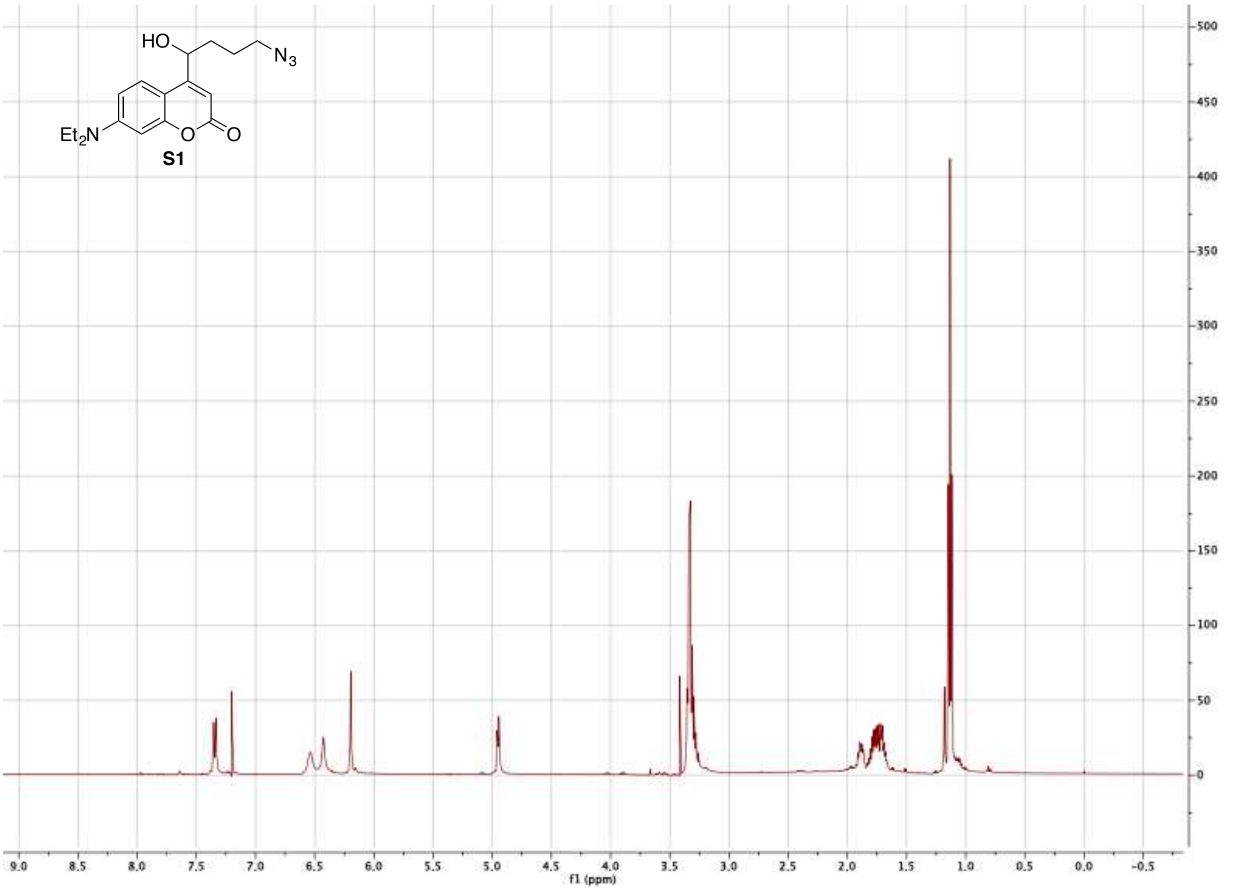
HO
N3
Et2N
O
O
S1
f1 (ppm)

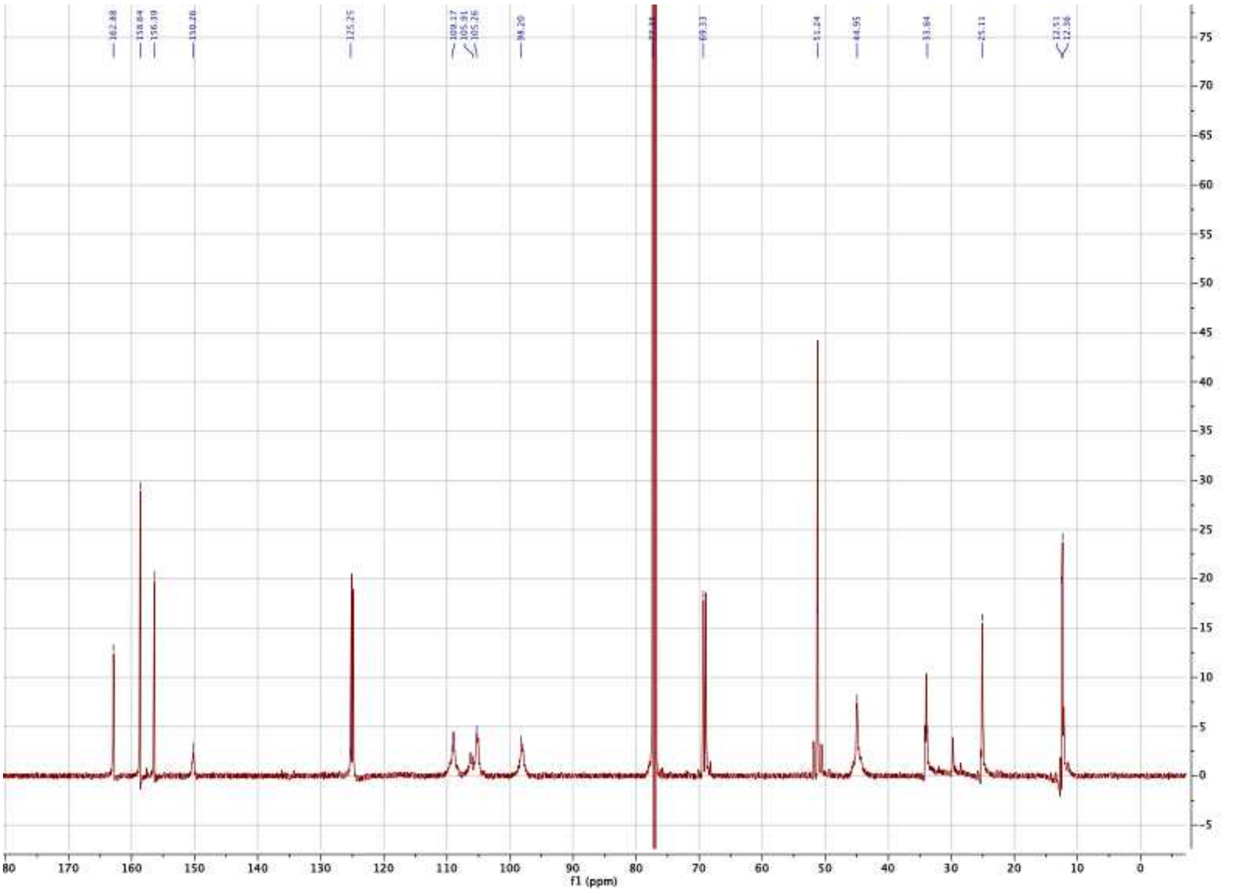
162.88
158.64
156.39
125.25
109.17
105.91
105.26
98.20
69.33
51.24
44.95
33.84
25.11
12.51
12.36
f1 (ppm)

#### Compound **(8)**

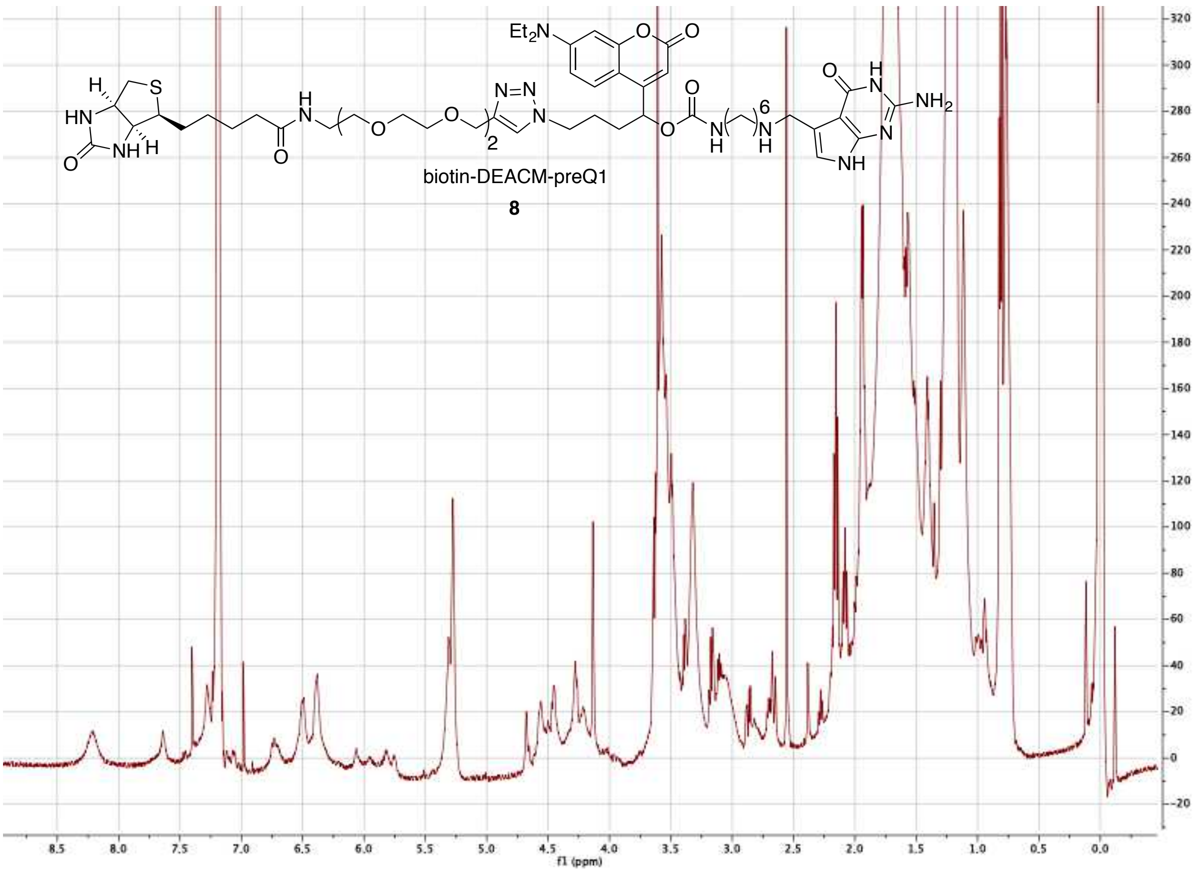

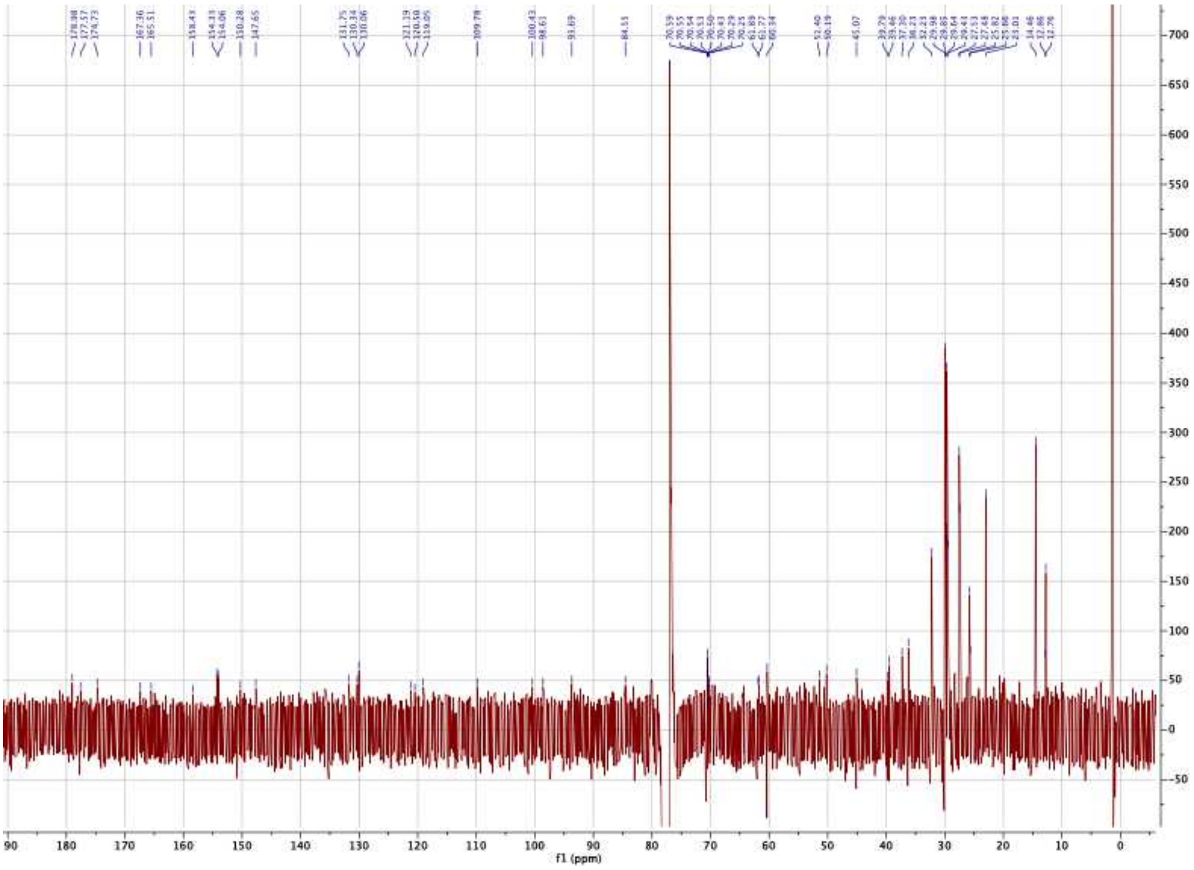

HRMS of compound **(8)**

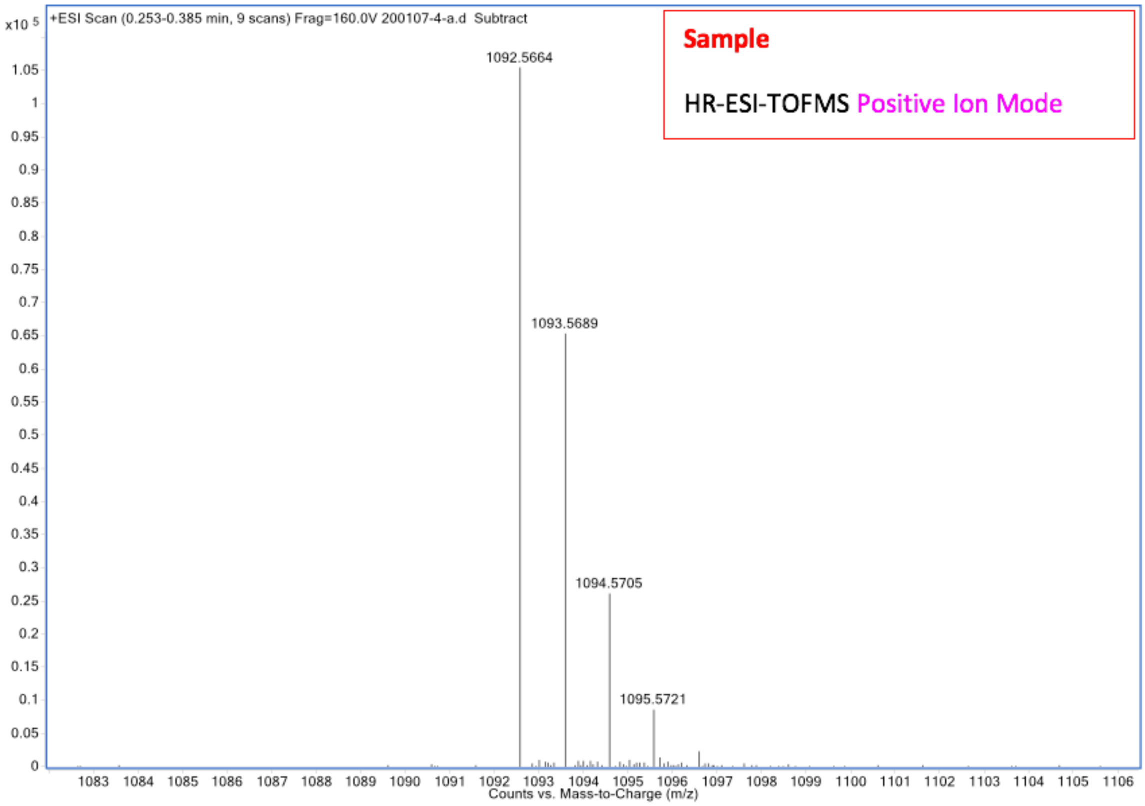

| Mass Measured | Theo. Mass | Delta (ppm) | Composition |
|---|---|---|---|
| 1092.5664 | 1092.5659 | -0.5 | $[C_{52}H_{78}N_{13}O_{11}S]^{+}$ |
